# Glutamine deficiency enhances nuclear localization of TCA cycle enzymes and epigenetic modifications, impairing myogenesis

**DOI:** 10.1101/2025.05.30.657066

**Authors:** Angad Yadav, Susan Schmitt, Wenxia Ma, James A. Mobley, Anna E. Thalacker-Mercer

**Affiliations:** University of Alabama at Birmingham, Birmingham, AL, USA

**Keywords:** Skeletal muscle metabolism, Myogenesis, TCA cycle, Succinylation, Glutamine metabolism, Chromatin accessibility

## Abstract

**Objective:** Skeletal muscle myogenesis, during development and regeneration, are primarily facilitated by the progeny of resident muscle stem cells, the muscle progenitor cells (MPCs). Intriguingly, extracellular glutamine (Gln) is required by MPCs. Disruptions in Gln availability or metabolism, impair MPC function and hinder skeletal muscle regeneration. Gln, a nutritionally non-essential amino acid (NEAA), which is most abundant in skeletal muscle, becomes conditionally essential during advancing age and catabolic states like sepsis, trauma, and strenuous situations. The intracellular mechanisms through which extracellular Gln availability affects MPC proliferation are poorly understood. Therefore, this study was aimed to examine how extracellular supply of Gln influences myogenesis through the spatial distribution of TCA-cycle enzymes.

**Methods:** Human primary muscle myoblasts (HSKM2) and murine MPCs (C2C12) were utilized for *in-vitro* experiments. To investigate the impact of Gln availability on MPC proliferation, proliferation assays, confocal imaging, succinyl-proteomics, and single-cell nuclei ATAC sequencing were used.

**Results:** Gln (2 mM) significantly enhanced MPC viability and cell cycle progression, while Gln depletion (0 mM) reduced cell number and induced G0/G1 arrest. Gln deficiency downregulated MyoD expression, impairing myogenic potential. Inhibiting intracellular Gln metabolism also limited proliferation, indicating the necessity of Gln catabolism for cell proliferation. Gln deficiency increased nuclear localization of TCA cycle enzymes (DLST, OGDH), elevated histone succinylation, and restricted chromatin accessibility of the myogenic regulatory factor, MyoD1, highlighting a Gln dependent metabolic-epigenetic nexus in myogenic regulation.

**Conclusion:** These findings demonstrate that extracellular Gln availability is a key regulator of myogenesis through its influence on metabolism, epigenetic modifications, and chromatin accessibility.

## INTRODUCTION

Skeletal muscle is composed of highly organized myofibers, formed by fusion of myocytes during development and regeneration through a coordinated myogenic process. To build or repair muscle, resident skeletal muscle specific stem cells (MuSC) and their progeny, muscle progenitor cells (MPCs), utilize an orchestrated myogenic process that includes MuSc activation and MPC proliferation^1^. Importantly, the myogenic process depends on the cellular microenvironment^2^. Disruptions to microenvironment, due to aging, nutrition, metabolic disorders, illness, and/or disease can impair MPC function and reduce myogenic potential^3^.

Extracellular nutrient availability, particularly nutritionally non-essential amino acids (NEAAs), plays a crucial role in regulating MPC function^4,5^. Among NEAAs, glutamine (Gln) is the most abundant in circulation and skeletal muscle^6^. Despite being primarily synthesized in skeletal muscle (endogenously), Gln becomes conditionally essential during catabolic states, such as sepsis, trauma, and injury, due to an increased demand^7^. Additionally, Gln depletion occurs during muscle remodeling processes associated with physiological and pathological stress, including inadequate recovery from overload training^8^, prolonged overtraining syndrome ^9^, and Duchenne muscular dystrophy (DMD)^10^. Studies have also demonstrated that extracellular Gln serves as a key metabolic substrate for rapidly proliferating cells^11^ including muscle cells^12^, further underscoring the importance of Gln for myogenesis; however, how Gln availability determines MPC function is unknown.

Gln is a key metabolic substrate that serves as an anaplerotic fuel for the tricarboxylic acid (TCA) cycle, supporting energy production, biosynthesis, and epigenetic regulation^13^. Intriguingly, emerging evidence suggests that under metabolic stress certain TCA cycle enzymes localize in the nucleus, where they influence epigenetic regulation by modifying chromatin accessibility^14–17^ and ultimately impairing proliferation^18^. The impact of Gln depletion on the subcellular distribution of TCA cycle enzymes and their subsequent impact on epigenetic modifications in MPCs have not been explored. This study investigates how Gln availability affects MPC proliferation through nuclear localization of TCA cycle enzymes (i.e., α-ketoglutarate dehydrogenase complex [αKGDHC] components—dihydrolipoamide succinyl transferase [DLST] and oxoglutarate dehydrogenase [OGDH]), histone succinylation, and chromatin accessibility of myogenic regulatory factors (MRFs). Our findings suggest that Gln depletion increases the nuclear localization of TCA cycle enzymes, leading to heightened histone succinylation and reduced chromatin accessibility at MRF regions. Targeting Gln metabolism to restore chromatin accessibility and enhance myogenic gene expression may provide a promising strategy for improving muscle regeneration and function in pathological states.

## MATERIALS AND METHODS

### Cell lines and media

Mouse MPC (C2C12) were obtained from the American Type Culture Collection (ATCC) and revived in pre-warmed *complete growth media* (Dulbecco’s Modified Eagle Medium [DMEM; 11960-044, Gibco], supplemented with 6 mM L-glutamine [Gln; 25030-081, Gibco], 1x Penicillin/Streptomycin [P/S; 15140-122, Gibco], and undialyzed 10% Fetal Bovine Serum [FBS; 35-011-CV, Corning, USA]). For the experiments, C2C12 cells were incubated in pre-warmed *customized growth media* (DMEM [11960-044, Gibco], 1x P/S, 10% dialyzed FBS, and various doses [0, 2, 6, and/or 12 mM] of Gln), as indicated in each experiment.

Human Embryonic Kidney 293 (HEK293) cells were cultured under similar conditions as C2C12 cells. HEK293 cells were grown in growth media for 96 hours without (0 mM) and with (2 mM) Gln supplementation, using the same customized DMEM-based media formulations as described above.

Human Skeletal Muscle Myoblast (HSKM2) primary cells were sourced from Lonza (CC-2580) and revived using Lonza’s recommended protocol, which includes Skeletal Muscle Basal Medium-2 (SKBM-2; CC-3246, Lonza) and Skeletal Muscle Growth Medium-2 (SKGM-2; CC-3244, Lonza).

For dialysis of the FBS (35-011-CV, Corning, USA), FBS was transferred into a dialysis bag with a 3.5 kDa cutoff membrane (VWR-28170-166, Spectrum Labs), securely knotted or clipped on both ends. The bag was submerged in 5 L of 1x Dulbecco’s Phosphate-Buffered Saline (DPBS, Gibco) inside a large container. A stir bar was placed in the container, and the entire setup was kept on a stir plate at 4°C, with continuous stirring maintained for 36 hours. The 1x DPBS buffer was replaced three times during the dialysis process. After dialysis, 2 g of charcoal (C-3345, Sigma) per 150 mL of FBS was added, and the solution was stirred at 4°C for 2 hours. The mixture was then sterilized by passing it through 0.2 µm filters inside a laminar air flow hood to remove the charcoal particles. Finally, the dialyzed and charcoal-treated FBS was aliquoted into 50 mL conical tubes and stored at -20°C until further use.

### Live and dead cell counting

Approximately 2,000 C2C12 cells were seeded per well in a 96-well culture plate and incubated in pre-warmed complete growth media at 37°C for 6 h to allow cell adhesion. The complete growth media was then replaced with customized growth media every 48 h, and cells were maintained for 5 days at 37°C, 5% CO_2_. For cell counting, the plate was incubated with pre-warmed media containing Hoechst 33342 (1:2000, Life Technologies) and propidium iodide (PI; 1:500, Thermo Fisher Scientific) at 37°C for 15 minutes. After staining, the media was replaced with fresh media, and cells were imaged using an automated imaging cytometer (ImageXpress PICO, Molecular Devices, USA). Hoechst stains all cell nuclei, including both live and dead cells, whereas PI only stains dead cells with compromised membranes. The number of live cells was calculated by subtracting the PI-positive cells from the total Hoechst-positive nuclei.

HEK293 cells were grown for 96 h and analyzed under the same conditions described above to serve as a non-muscle cell model. This was done to confirm whether Gln availability impacts cell proliferation similarly in non-muscle cells.

### Cell viability assessment

To validate C2C12 cell viability, an MTS assay (CellTiter 96® AQueous One Solution Cell Proliferation Assay, Promega, USA) was conducted. The reagent, which contains MTS (a tetrazolium compound) and PES (an electron coupling reagent), was added directly to the C2C12 cells in the 96-well culture plate. After a 3 hours incubation, absorbance was measured at 490 nm using a microplate reader (Bio-Rad, USA), with absorbance values correlating to the number of viable cells.

### Pharmacological inhibition of glutaminase

To investigate the role of Gln catabolism in MPC proliferation, the pharmacological inhibitor CB-839 (22038, Cayman, USA)^19^ was used, which inhibits glutaminase (GLS), the enzyme responsible for converting Gln to glutamate (Glu). A 10 µM solution of CB-839 was prepared using dimethyl sulfoxide (DMSO; Thermo Fisher Scientific) as the vehicle. C2C12 cells were treated with three different conditions: (i) customized growth media containing 2 mM Gln, (ii) growth media containing 2 mM Gln and DMSO, and (iii) growth media containing 2 mM Gln and 10 µM CB-839. Similarly, the effect of CB-839 on HSKM2 cells was assessed using Lonza-recommended media containing 2 mM Gln, with or without DMSO and CB-839. To evaluate the impact of CB-839 on cell proliferation, cell counting was performed every 48 hours using an automated imaging cytometer, as described previously.

### Quantitative RT-PCR

C2C12 cells were seeded at 15% confluency on 60 mm culture plates for 5 days in customized growth media containing various doses (0, 2, 6, and 12 mM) of Gln. Media was aspirated, and 1 mL of TRIzol (5596018, Invitrogen) was added to each plate. The cells were scraped, transferred to a 1.5 mL Eppendorf tube, and left at room temperature (RT) for 5 minutes. Bromochloropropane (BCP) was then added at a 1:5 ratio with Trizol, the tube was shaken vigorously for 1 minute and left for another 5 minutes at RT. The mixture was centrifuged at 12,000x g at 4°C for 15 minutes. The clear aqueous top layer was carefully removed to avoid DNA and protein contamination and transferred to a new tube. Next, isopropanol was added in 1:1 ratio of top layer, mixed gently by inverting the tube 4-6 times, and left to sit for 10 minutes at RT. The mixture was centrifuged at 12,000× g at 4°C for 10 minutes. The isopropanol was poured off, and 700 µL of 75% ethanol in diethyl pyrocarbonate (DEPC)-water was added. The tube was decanted, and the pellet was washed three times with 700 µL of 75% ethanol in DEPC-water, gently shaken, and centrifuged at 7500× g at 4°C for 5 minutes each time. After the final wash, the samples were centrifuged at 7500× g at 4°C for 1 minute. The ethanol was poured off, and the pellets were air-dried without over-drying at RT. Then, 50 µL of DEPC-water was added to the RNA pellet and left for 10 minutes to dissolve completely. The RNA concentration was measured spectrophotometrically. cDNA was prepared by using manufacture’s protocol of high-capacity RNA-to-cDNA kit (4368814, Thermo scientific). SYBR green (A25742, Thermo scientific) method was used to measure the level of gene expression on QuantStudio-3 (Thermo Fisher). The expression of PAX7 and MyoD genes were normalized with the β-actin expression. The primer sequences for PAX7, MyoD and β-actin are listed in **Table S1**.

### In-cell western

Using previously developed methods^4^, in-cell western assay was performed to quantify the protein levels of the myogenic regulatory factors-PAX7 and MyoD during proliferation in a 96-well culture plate. To achieve this, the 96-well plate was first coated with 1% collagen (1:100 in 0.02 M acetic acid) and washed two times with 1× DPBS before seeding cells. The C2C12 cells were incubated in customized growth media with (2 mM) and without (0 mM) Gln for 5 days at 37°C, 5% CO2. Cells were then fixed with 4% paraformaldehyde (PFA) for 10 minutes at RT, washed five times with 1× DPBS, and permeabilized with 0.2% Tween-20 (T20) for 10 minutes at RT. After three washes with 1× DPBS, blocking was performed using 1% bovine serum albumin (BSA) in 1× DPBS for 1 hour at RT. Primary antibodies against MyoD and PAX7 (**Table S2**) were diluted in blocking solution and incubated overnight at 4°C. After three washes with 1× DPBS containing 0.1% T20 (DPBST), secondary antibodies (**Table S2**) were diluted 1:800 in blocking buffer and added along with a cell normalization stain (Cell Tag-520, diluted 1:1000 in blocking buffer). The plate was incubated for 1 h in the dark at RT, followed by washed with 1× DPBST for 5 times. The intensity of stained proteins and cell tags was measured using the Odyssey Imaging System (LI-COR). Protein levels (680 or 800 intensity) were normalized to Cell Tag-520 intensity using Image Studio 6.0 software.

### FACS analyses

For cell cycle analyses, the C2C12 cells grown for 5 days in customized growth media with (2 mM) and without (0 mM) Gln, were pelleted. We have followed the protocol developed in our laboratory^4^.

### Western blot analyses

C2C12 cells were seeded at approximately 15% confluency in 150 mm culture plates and cultured in customized growth media with varying Gln concentrations (0, 2, 6, and 12 mM) for 5 days, as described previously.

For whole cell lysate extraction, cells were washed with 1× DPBS (Gibco) and lysed using 1× cell lysis Radioimmunoprecipitation Assay (RIPA) buffer (50mM Tris-HCl [pH 7.4-7.6], 150 mM NaCl, 1% NP-40, 0.5% sodium deoxycholate, and 0.1% Sodium dodecyl sulphate [SDS]) containing 1× Halt cocktail inhibitor (1861280, Thermo Scientific). The lysate was kept on ice for 10 minutes, after which cells were scraped using a cell scraper and transferred into 1.5 mL Eppendorf tubes. The lysate was incubated on ice for 30 minutes, with vortexing every 10 minutes for 15 seconds at high speed. The lysate was then centrifuged at 20,000 × g for 25 minutes, and the supernatant was collected.

For nuclei lysate extraction, cells were harvested on day-5 using TrypLE (12605-028, Gibco) followed by centrifugation at 400× g for 5 minutes, and nuclei were isolated using the Nuclei Pure Prep Kit (NUC-201, Sigma). The quality of nuclei purification was checked using nuclei staining dye on hemocytometer. The purified nuclei pellet was lysed in RIPA buffer containing 1× Halt inhibitor and incubated on ice for 30 minutes, with vortexing every 10 minutes for 15 seconds at high speed. After centrifugation at 20,000× g for 25 minutes, the supernatant was collected as the nuclear lysate. Protein concentration was determined using the BCA assay (23227, Thermo Scientific). Protein loading samples were prepared by mixing lysates with Laemmli buffer (1610747, Bio-Rad) containing 10% β-mercaptoethanol (βME) and heating at 95°C for 10 minutes.

A total of 15 μg of protein per sample was resolved on 4–20% SDS-PAGE gels (Bio-Rad, USA) and transferred onto PVDF membranes (Bio-Rad, USA). Membranes were blocked with 5% skimmed milk (M17200, RPI Corp., USA) in 1× Tris-buffered saline with Tween-20 (TBST) for 1 h at RT with gentle shaking (22–25 RPM), followed by three washes with 1× TBST (5 minutes each). Membranes were then incubated overnight at 4°C with primary antibodies against MyoD, MyoG, SLC1A5, DLST, OGDH, KAT2A, SIRT1, SIRT7, pan-lysine succinylation, α-tubulin, and GAPDH (**Table S2**). After three washes with 1× TBST (5 minutes each), membranes were incubated with secondary antibodies (**Table S2**) for 1 h at RT. Following another set of washes, membranes were treated with chemiluminescence substrate (Sativa, Global Life Sciences, UK) and immediately visualized using a ChemiDoc system (Bio-Rad). Protein expression was normalized to housekeeping proteins α-tubulin, GAPDH, and total protein. Quantification of protein bands was performed using ImageJ software.

Similarly, HEK293 cells cultured under the same Gln (0 vs 2 mM) conditions for 96 h and were analyzed for DLST, α-tubulin, and total protein to compare the impact of Gln availability on non-muscle cell protein levels.

### Immunofluorescence

C2C12 cells were seeded at approximately 15% confluency on 12 mm coverslips coated with 1% collagen (1:100 dilution in 0.2 mM acetic acid) in 24-well culture plates. The cells were then incubated at 37°C in customized growth media with (2 mM) and without (0 mM) Gln for 5 days. Cells were stained with MitoTracker (250 nM, Thermo Fisher) for 30 minutes at 37°C, washed with phenol-free basal DMEM (Gibco), and fixed with 4% PFA for 15 minutes at RT. Following four washes with 1× DPBS, cells were permeabilized with 0.1% Triton X-100 for 10 minutes and then washed five times with 1× DPBS. Cells were blocked with 10% FBS in 1× DPBS for 30 minutes at RT. After blocking, the cells were incubated overnight at 4°C with primary antibodies against DLST and Lamin-B1 (**Table S2**). The next day, the cells were washed five times with 1× DPBS and incubated with secondary antibodies (**Table S2**) for 1 h at RT. Following secondary antibody incubation, the cells were washed five times with 1× DPBS. Nuclear staining was performed using Hoechst (1:2000) for 15 minutes at RT, followed by a final 5-minute wash with DPBS. Coverslips were mounted using ProLong Antifade Mountant (Thermo Scientific) and sealed with clear nail polish. The intracellular spatial distribution of DLST was visualized using a confocal microscope (Nikon-1XR, Nikon Instruments) at 60× magnification with a Z-stack acquisition.

### Quantification of total succinyl-proteomics

The whole cell lysate collected from C2C12 cells grown in customized growth media with (2 mM) and without (0 mM) Gln for 5 days. 400 µL of the whole cell lysate was divided into two separate tubes: one incubated with pan anti-succinylation antibody (1:500; **Table S2**), and the other with untagged Anti-mouse IgG (**Table S2**) for overnight at 4°C. Magnetic beads (88802, Thermo Scientific) were added to the lysate mixture at a 1:10 ratio. The preservative solution from the beads was removed using a magnetic rack (Millipore), and the beads were washed once with 1000 µL of washing buffer (150 mM NaCl, 1 mM EDTA, 1% Triton X-100, 10 mM Tris, pH 7.4, Milli-Q water). The antigen-antibody complex was incubated with the pre-washed beads while rotating at RT for 20 minutes. After incubation, the beads were separated using a magnetic rack and washed five times with the same washing buffer. After the final wash, the bead-bound complex was resuspended in 50 µL of protein loading buffer (4× Laemmli buffer with 10% βME) and heated at 95°C for 10 minutes. The supernatant was collected by centrifugation at 500x g for 5 minutes at 4°C, loaded onto an SDS-PAGE gel, and subjected to western blot analysis.

Whole cell lysate sample, immunoprecipitated with anti-pan lysine-succinylation, was subsequently analyzed for succinyl-proteomics using untargeted approach. The entire eluate was reduced with dithiothreitol (DTT) and denatured at 70°C for 35 minutes before loading onto a 10% Bis-Tris protein gel for separation. After staining the gel overnight with colloidal Coomassie for visualization, the lane was divided into three molecular weight fractions based on experimental optimization. Each fraction was equilibrated in 100 mM ammonium bicarbonate and digested overnight with Trypsin Gold, Mass Spectrometry Grade (V5280, Promega) according to the manufacturer’s instructions. The peptide extracts were then reconstituted in 0.1% formic acid/ddH_2_O at a concentration of 0.1 µg/µL. Mass spectrometry was performed, and the resulting data were processed, searched, filtered, grouped, and quantified as described in a previous study^20^.

### Single nuclei ATAC sequencing

C2C12 cells were cultured in customized growth media with (2 mM) and without (0 mM) Gln for 5 days. Following incubation, cells were trypsinized using TrypLE, collected in 15 mL Falcon tube, and centrifuged at 300× g for 5 minutes at 4°C. The cell pellet was resuspended in 1 mL DPBS containing 0.04% BSA and transferred to a 2 mL microcentrifuge tube. The Falcon tube was rinsed with 0.5 mL of DPBS containing 0.04% BSA, and the rinse was added to the cell suspension in the 2 mL microcentrifuge tube. Cells were centrifuged again at 300× g for 5 minutes. After removing the supernatant, the pellet was resuspended in 1 mL DPBS + 0.04% BSA, filtered through a 40 µm Flowmi Cell Strainer, and the cell concentration was determined using a Countess Automated Cell Counter. For nuclei isolation and single-cell ATAC sequencing, the 10x Genomics protocol (CG000169) was followed. Briefly, ∼10,000 nuclei were processed per sample, subsequent library preparation and high-throughput sequencing using the Chromium Next GEM Single Cell ATAC Reagent Kits v2 (CG000496). The Cell Ranger ATAC count tool was used to analyze supplemented (2 mM) and depleted (0 mM) ATAC-seq samples. FASTQ files were processed using a reference genome (mm10 Reference: 2020-A-2.0.0) pre-downloaded from 10x Genomics. For each sample, the cellranger-atac count pipeline was run separately, generating barcode-by-peak matrices, annotated peak files, and summary outputs. The resulting data, including filtered peak matrices and Loupe Browser files, were used for downstream analyses such as differential chromatin accessibility between control verses treatment.

### Statistical analysis

To compare the effect of Gln dose (with and without Gln) on cell cycle analysis, enzyme assays, and quantitative assays, we used unpaired two-tailed t-tests. For multiple comparisons in immunoblots and cell counting experiments involving various doses of Gln (0, 2, 6, and 12 mM) and GLS inhibitor treatment (control, vehicle and inhibitor), one-way ANOVA followed by Tukey’s post hoc test was employed. Statistical analyses were conducted using GraphPad (version-10), with a significance level set at p < 0.05.

## RESULTS

### Extracellular Gln is essential for MPC proliferation

We compared the effect of various concentrations of Gln (0, 2, 6, 12 and 19 mM) on MPC proliferation (**Figure 1A**) and death (**Figure 1B**). Despite the higher concentration of Gln previously identified in skeletal muscle (i.e., 19 mM)^6^, 19 mM Gln resulted in enhanced MPC death (data not shown) and thus, was removed from subsequent studies. We identified that 2 mM (vs. 0, 6, and 12 mM) extracellular Gln resulted in the significantly higher MPC number over 5 days compared to other concentrations. Conversely, Gln depletion (0 mM) retarded cell number. An MTS assay confirmed these results; the MTS cell viability assay demonstrated significantly higher MPC viability in presence of 2 mM Gln as compared to 0 mM Gln (**Figure 1C**). These findings are consistent with observations in other proliferating cell types, including HEK293 cells, where extracellular Gln availability similarly regulated cell number and viability (**Figure S1**).

**Figure 1.**
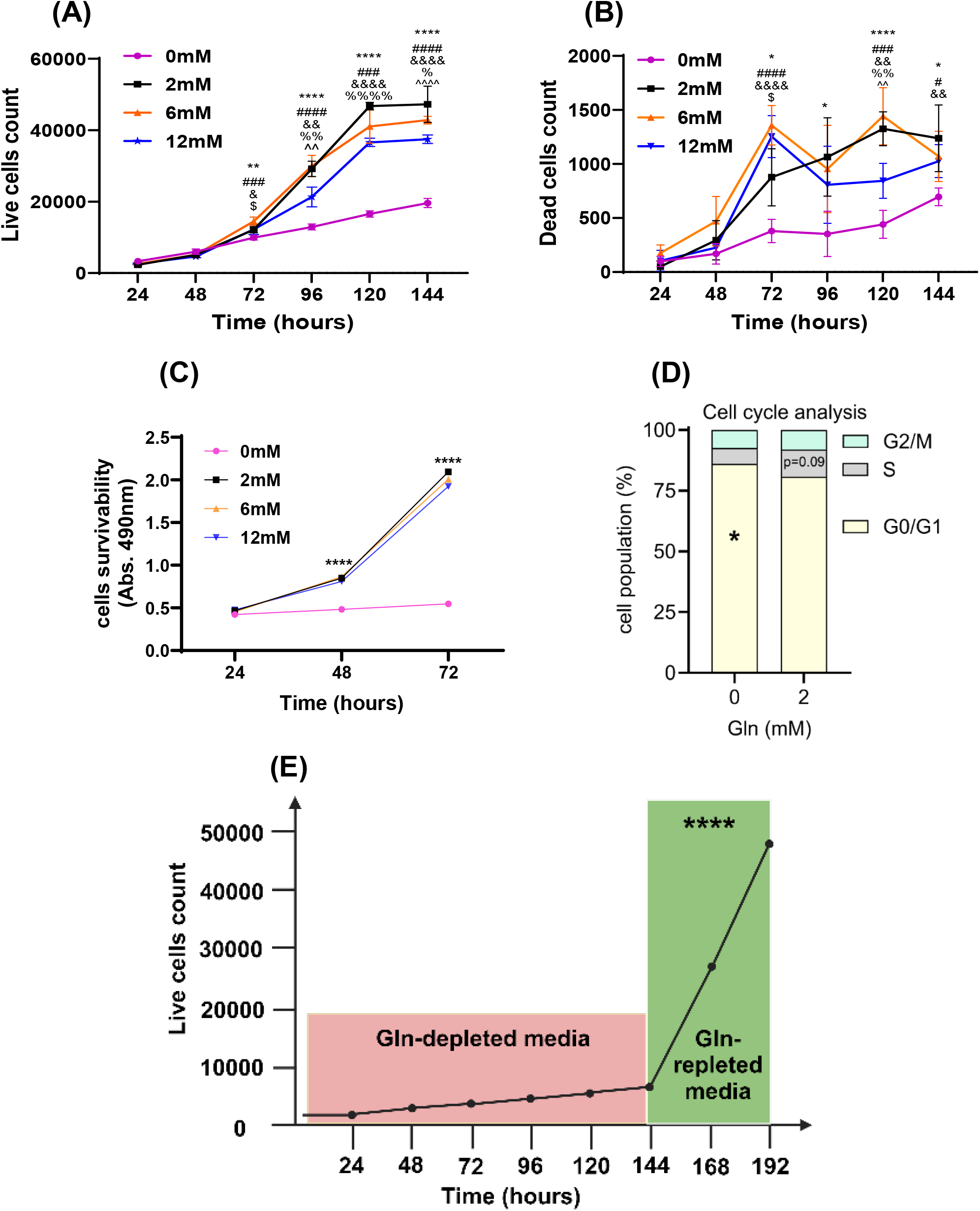
Impact of extracellular Gln availability on MPC proliferation. **(A)** Live and **(B)** dead cells count of C2C12 cells grown for 5 days with (2, 6 and12 mM) and without (0 mM) of Gln in growth media, n=6. (^**\***^ indicates the significant difference between 0 mM glutamine vs 2 mM Gln; **#** indicates the significant difference between 0 mM Gln vs 6 mM Gln; & indicates the significant difference between 0 mM Gln vs 12 mM Gln; **$** indicates the significant difference between 2 mM Gln vs 6 mM Gln; **%** indicates the significant difference between 2 mM Gln vs 12 mM Gln; **^** indicates the significant difference between 6 mM Gln vs 12 mM Gln). Cells were counted by using co-stained with propidium iodide (dead cells) and Hoechst-33342 (cell nuclei) on imaging cytometer. The difference of propidium iodide and Hoechst indicated the total number of live cells. **(C)** MTS assay for validating the impact of extracellular supply of Gln on cell viability. (^****^indicates the significant difference between 0 mM vs others Gln supplemented media. **(D)** Effect of Gln availability on cell cycle distribution. Cells in Gln -depleted conditions showed increased arrest in the G0/G1 phase, n=3. **(E)** Effect of switching C2C12 cells from glutamine-depleted to glutamine-repleted media for 48 hours on cell numbers, n=6. Cells returned to a normal proliferative condition upon switching from depleted to repleted media. (^****^indicates the significant difference in cell number between Gln depleted vs repleted media.

Gln availability has been implicated in cell cycle regulation, as observed in non-muscle cell lines^21^. In our study, we investigated the impact of extracellular Gln on cell cycle progression. Gln depletion impaired cell cycle progression, with MPCs more likely to arrest in the G0/G1 phase (**Figure 1D**). To determine whether this arrest was reversible, C2C12 cells were cultured in Gln-depleted growth media for five days, leading to a significant reduction in proliferation. When switched to replete growth media, containing 2 mM Gln for 48 hours, MPCs regained their proliferative capacity, as indicated by an increase in cell number (**Figure 1E**).

Proliferation and differentiation of MPCs requires the coordinated activity of MRFs such as paired box transcription factor 7 (Pax7)^22^, myoblast determination protein 1 (MyoD)^23^, and myogenin (MyoG)^24^. To further assess the effects of Gln deficiency on the myogenic process, we examined the mRNA and protein levels of Pax7, MyoD, and MyoG, in cells cultured with and without Gln. Our analysis revealed that 0 mM Gln (vs. 2, 6 and 12 mM Gln) significantly decreased the mRNA and protein levels of Pax7 and MyoD (**Figure 2A-D**). Given that PAX7 and MyoD are critical for MPC proliferation and myogenesis, their reduced expression suggests that Gln deficiency impairs myogenic potential. Additionally, we assessed MyoG to determine if the cells differentiated in 0 mM Gln but observed no difference in MyoG protein levels between 0 and 2 mM Gln (**Figure 2E**). These results support a prior study^12^, which demonstrated that extracellular Gln is imported by MuSCs and promotes cell proliferation and differentiation. While our observations partially contrast a study on low-birth-weight piglets, where oral Gln supplementation supported proliferation but did not significantly alter Pax7 or MyoD mRNA levels, the expression of MRFs in whole muscle homogenates, from the piglets, might be too small to detect^11^.

**Figure 2.**
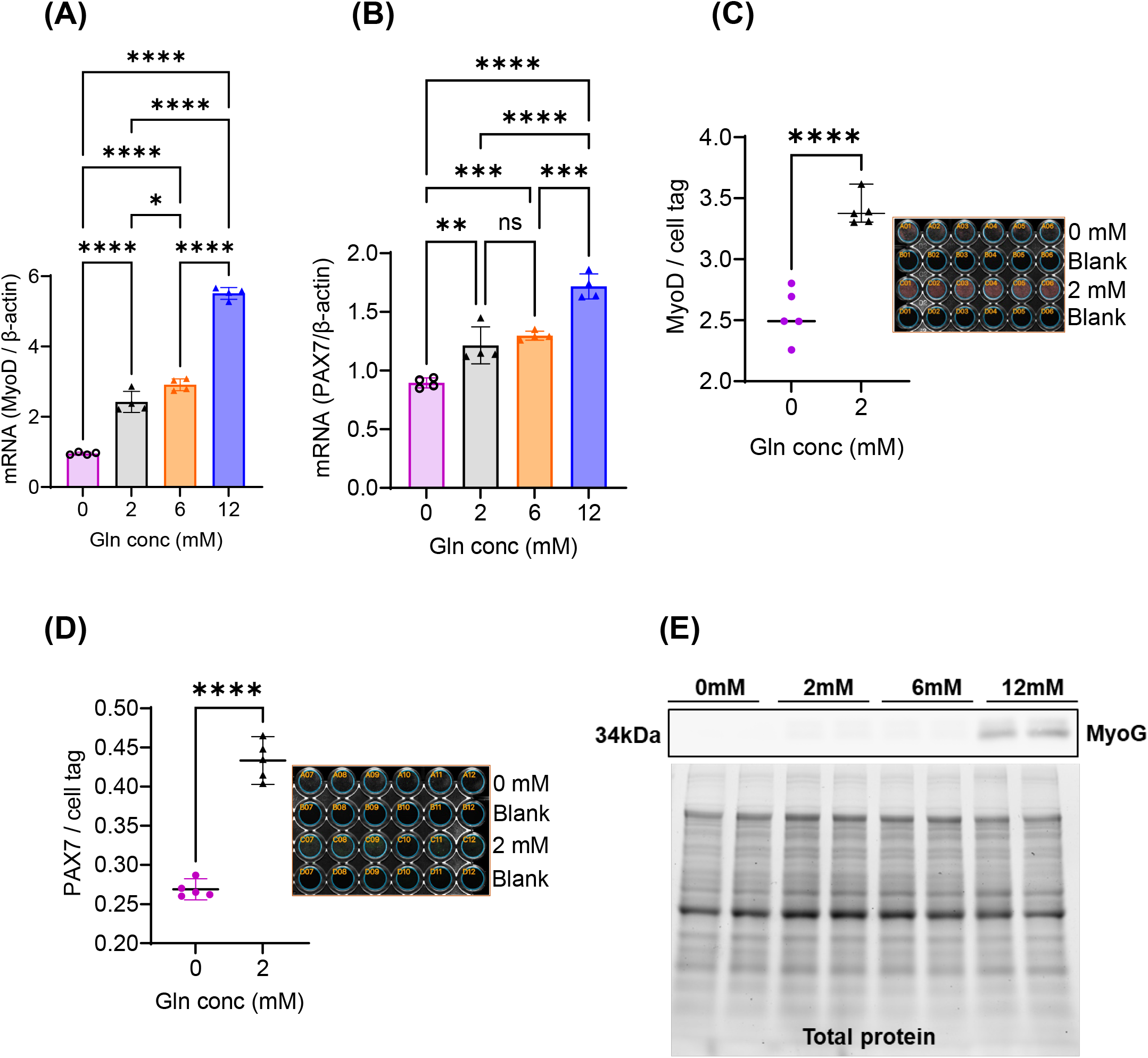
Extracellular availability supports myogenic regulatory factors. mRNA levels of PAX7 **(A)** and MyoD **(B**) was decreased in Gln deficient condition; n=5. Protein levels of PAX7 **(C)** and MyoD **(D)** was decreased in Gln deficient condition; n=3. **(E)** Representative immunoblot of MyoG showed that cells are not in differentiation state in 0 vs 2 mM Gln; n=2. Protein levels of MyoD and PAX7 were quantified using the In-cell Western technique with Li-COR. ^*^ p<0.05; ^**^ p<0.01; ^***^ p<0.001; ^****^ p<0.0001; ns-non significant.

### Inhibiting intracellular Gln catabolism restricts MPC proliferation

The transport of Gln into cells is facilitated by the transporter SLC1A5 (**Figure 3A**). The Gln transporter, mitochondrial variant, is SLC1A5_var^25^. Western blot analysis showed a significant increase in both transporters under Gln-depleted conditions (**Figure 3B**), indicating an adaptive response to support cell survival. Once in the mitochondria Gln is catabolized to Glu (via glutaminase [GLS]) which enters the TCA through further metabolism to α-KG. Despite elevated protein levels of Gln transporters, we observed decreased GLS catabolic activity (**Figure 3C**) and subsequently reduced α-KG (**Figure 3D**) level, with depleted Gln availability. To determine whether the intracellular Gln metabolism is essential for MPC proliferation, we inhibited GLS using CB-839, a potent GLS inhibitor (GLSi)^19^. In both C2C12 and HSKM2 cells, live cell counts significantly decreased with GLSi compared to control (**Figures 3E** and **3G**). Concurrently, there was a decline in dead cell number with GLSi for both cell types (**Figures 3F and 3H**). These results suggest that Gln metabolism is necessary for MPC proliferation.

**Figure 3.**
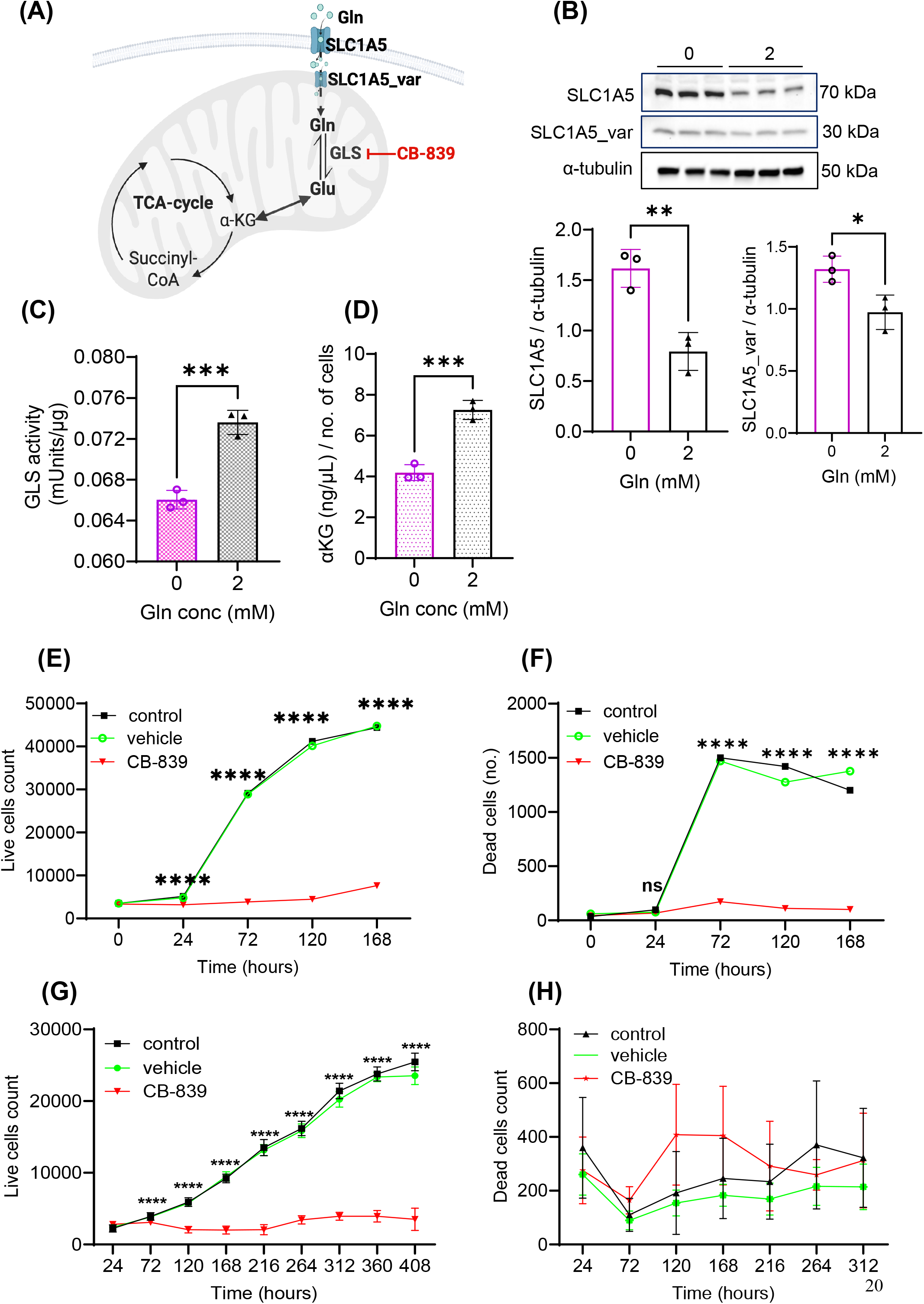
Effect of glutamine availability on Gln-transporters and subsequent downstream metabolism. **(A)** Schematic of the potential mechanism by which extracellular Gln entered the TCA cycle and the action of the GLS inhibitor (CB-839). **(B)** Immunoblot showed that the protein levels of Gln transporters were elevated in Gln-deficient conditions, n = 3. **(C)** Enzymatic assay showed that GLS activity was lowered in Gln-deficient conditions. **(D)** Quantification assay of α-KG in cell lysates from cells grown for 5 days in growth media with and without Gln, n = 3. **(E)** Live and **(F)** dead C2C12 cell counts after treatment with CB-839 for 5 days in growth media, n = 6. **(G)** Live and **(H)** dead HSKM2 primary cell counts after treatment with GLS inhibitor for 408 hours in proliferation media, n = 6. p < 0.05; ** p < 0.01; *** p < 0.001; **** p < 0.0001; ns – not significant.

### Gln availability regulates TCA cycle enzymes and subsequently protein succinylation

α-KG, a metabolic product of Gln, can be metabolized through both reductive TCA (r-TCA) and oxidative TCA (o-TCA) cycle enzymatic reactions. Products of these reactions include acyl-CoAs (e.g., acetyl- and succinyl-CoA) that can be used not only as metabolic substrates, but also as acyl donors for protein lysine acylations, a type of post-translational protein modification. Protein acylation is a reversible lysine residue modification that impacts protein localization, structure, and activity ultimately influencing metabolism and other cellular processes such as transcriptional regulation^26^. We reasoned that Gln availability modifies cell state through acyl modifications, either acetylation or succinylation^27,28^ (**Figure 4A**). In r-TCA, α-KG availability can impact acetyl-CoA availability, yet we did not observe a significant difference in acetyl-CoA levels regardless of Gln concentration (**Figure S2A**). Similarly, SIRT1, a sirtuin protein involved in deacetylation, exhibited no significant difference between 0 mM and 2 mM Gln at whole-cell (**Figure S2B**) and subcellular levels (**Figure S2C**), indicating that Gln deficiency does not appear to impact acetyl-CoA.

**Figure 4.**
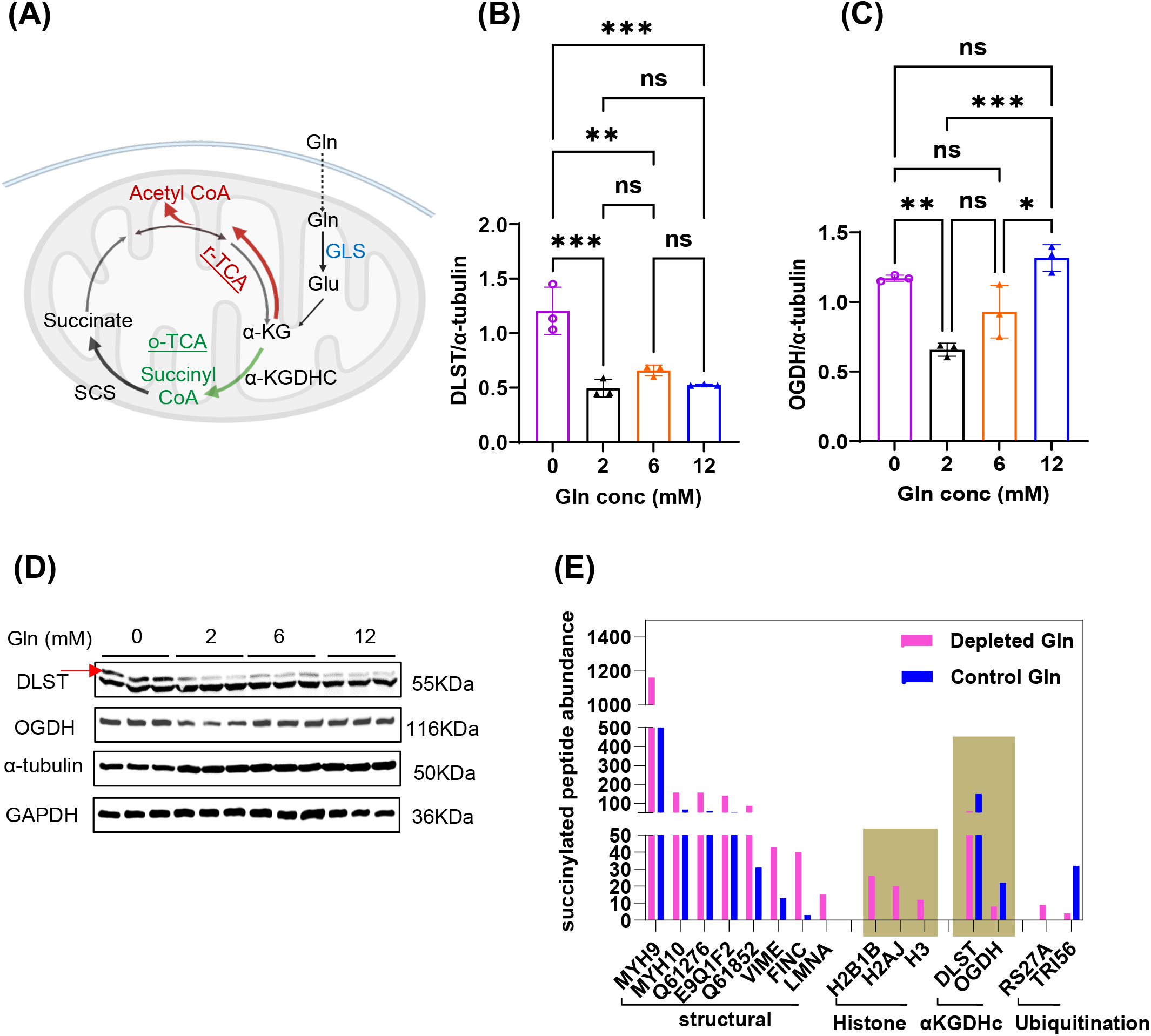
Glutamine availability favors the oxidative mode of TCA cycle and regulate pan-succinylation. **(A)** Schematic illustrating the influence of extracellular supply on the oxidative and reductive modes of the TCA cycle. o-TCA: oxidative TCA; r-TCA: oxidative TCA **(B)** Normalized DLST protein levels decreased with increasing Gln availability (n = 3). **(C)** Normalized OGDH protein levels showed variability in response to different Gln concentrations (n = 3). **(D)** Western blot images of DLST and OGDH under varying Gln concentrations (n = 3 per condition). **(E)** Top-most differentially succinylated proteins identified through succinyl-proteomics showed increased succinylation in Gln-deficient condition. ^*^ p<0.05; ^**^ p<0.01; ^***^ p<0.001; ^****^ p<0.0001; ns-non significant.

We hypothesized that intracellular Gln availability, during proliferation, might be signaled through o-TCA cycle enzyme regulation. To investigate the downstream fate of α-KG, through o-TCA cycle, we examined components of the α-KG dehydrogenase complex (α-KGDHC). α-KGDHC components, DLST and OGDH, total protein levels were significantly elevated under Gln-depleted conditions (**Figure 4B-D**), suggesting enhanced succinyl-CoA synthesis and potentially changes in succinyl protein modifications. Notably, a similar increase in DLST protein levels under Gln-deficient conditions was observed in HEK293 cells, supporting the idea that Gln availability broadly regulates α-KGDHC enzyme levels across proliferating cell types (**Figure S1**). Next, we examined the impact of Gln availability on global protein succinylation via mass spectrometry. We identified 87 differentially (0 vs. 6 mM) succinylated proteins (**Table S3**), with the 15 most abundant proteins belonging to cytoskeletal proteins, histones, TCA cycle enzymes, and ubiquitination-related proteins. Among these, cytoskeletal proteins and histones showed increased succinylation, while TCA cycle enzymes and ubiquitination-related proteins showed decreased succinylation (**Figure 4E**). Intriguingly, under Gln-deficient conditions, we observed a decrease in succinylation of DLST and OGDH. Recognizing that changes in protein modifications can influence protein localization, we questioned whether α-KGDHC enzymes were localizing outside the mitochondria in Gln deficient conditions.

### Gln deficiency increases nuclear localization of TCA cycle enzymes

Beyond its established biosynthetic and bioenergetic roles in mitochondria, studies conducted in non-muscle tissues have shown that components of the TCA cycle also localize in the nucleus, where they play a crucial role in regulating developmental processes through epigenetic modifications^14–17,29^. Under Gln-depleted conditions, we observed increased nuclear localization of DLST (**Figure 5A-B**) and OGDH (**Figure 5C**), suggesting a potential increase in nuclear protein succinylation. These components of the nuclear α-KGDHC interact with KAT2A, a lysine succinytransferase that enzymatically transfers succinyl groups to lysine residues of nuclear proteins (Wang et al., 2017). Correspondingly, we observed elevated KAT2A levels in the nucleus (**Figures 5D**) and no change in protein levels for nuclear deacylases, SIRT7 (**Figure 5E**). The immunoblot images of DLST, OGDH, KAT2A, and SIRT7 are shown in **Figure 5F**, and the image of total proteins is shown in **Figure 5G** These findings support the increase in histone succinylation we observed in our succinylation proteomics data, increased histone succinylation (i.e., histones [H2B1B and H2AJ and H3], **Figure 4G**) in Gln-deficient conditions. These findings indicate that Gln-deficiency increases nuclear localization of α-KGDHC and subsequently increases histone succinylation, which could ultimately impact chromatin structure.

**Figure 5.**
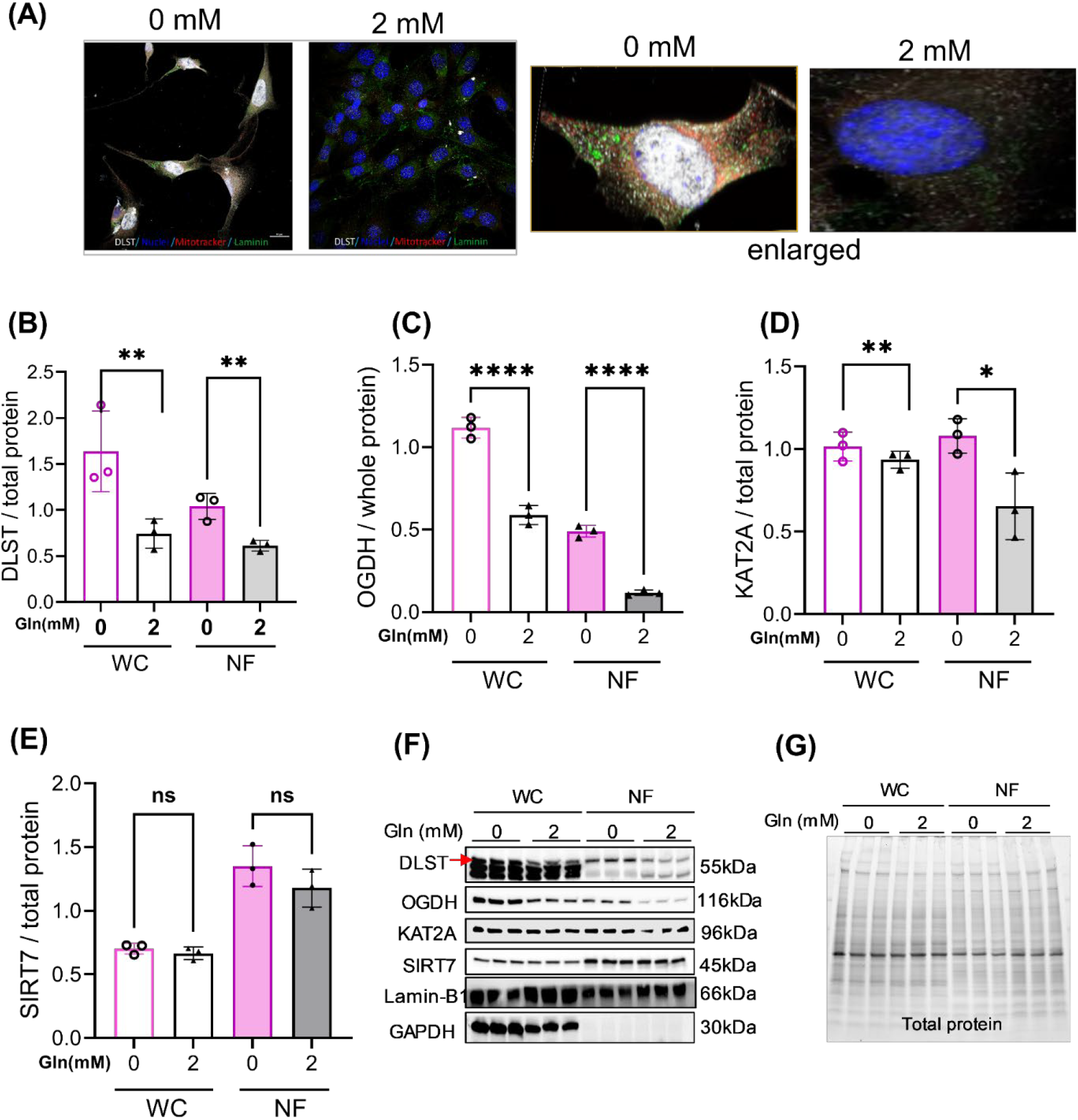
Glutamine deficiency induces nuclear localization of intermediary metabolic mediators. **(A)** Confocal microscopy images showing increased nuclear localization of DLST in the cells treated with 0 mM vs 2 mM Gln. Images represent maximum intensity projections of z-stacks (60× magnification) of C2C12 cells cultured for 5 days in a growth media. Scale bar = 20 µm. Cells were stained for DLST (white), MitoTracker (red), laminin (green), and DAPI (blue). **(B)** Normalized DLST protein levels in whole-cell and nuclear fractions decreased with increasing Gln availability (n = 3). **(C)** Normalized OGDH protein levels in whole-cell and nuclear fractions decreased with increasing Gln availability (n = 3). **(D)** Normalized KAT2A protein levels in whole-cell and nuclear fractions decreased with increasing Gln availability (n = 3). **(E)** SIRT7 protein levels remained unchanged in both whole-cell and nuclear fractions across Gln concentrations (n = 3). **(F)** Immunoblots of DLST, OGDH, KAT2A and SIRT7 for whole cell and nuclear fraction. **(G)** Immunoblot of total protein of whole cell and nuclear protein from cells treated with 0 mM vs 2 mM Gln. ^*^ p<0.05; ^**^ p<0.01; ^***^ p<0.001; ^****^ p<0.0001; ns - non significant.

### Elevated histone succinylation restricts chromatin accessibility in myogenic regulatory factor regions

We hypothesized that Gln deficiency alters the chromatin landscape and thereby impacts gene expression related to myogenesis. To explore how extracellular supply of Gln affects chromatin accessibility, single nuclei ATAC sequencing was performed on C2C12 cells grown in customized growth media with (2 mM) and without (0 mM) Gln for 5 days. Noticing reduced expression of MRF proteins and heightened succinylated peptide abundance of histone proteins in Gln-deficient cells, the chromatin accessibility region of MyoD1 gene was less accessible in depleted (0 mM) condition (**Figure 6A**) as compared to supplemented (2 mM) condition (**Figure 6B**), suggesting a restriction in chromatin structure that may hinder gene expression of MyoD1 in the Gln-deficient condition. Indeed, in earlier studies we demonstrated reduced mRNA levels for MyoD (**Figure 2B**). Previous studies showed that histone succinylation specific modifications such as H2BK34^30^, H3K79^29^, H3K122^26^, H4K77^31^ alters nucleosome unwrapping rates and impacts DNA accessibility, in non-muscle cells. Additionally, motif enrichment analysis, which identifies short, recurring patterns in DNA that play critical roles in gene regulation, demonstrated that MRFs (myf5, myf6, myoD1 and myoG) were significantly enriched in the Gln supplemented (2 mM) condition (**Figure S3**). Collectively our data demonstrate that Gln availability mediated histone succinylation is a regulator of chromatin structure and gene expression in myogenesis. These results suggest that extracellular Gln availability is crucial for regulating chromatin accessibility, particularly for genes essential for MPC proliferation, such as MyoD, and is accompanied by an increase in succinylation of histones and structural proteins. These findings collectively highlight the significance of Gln availability in differential succinylation of proteins involved in regulating MPC proliferation.

**Figure 6.**
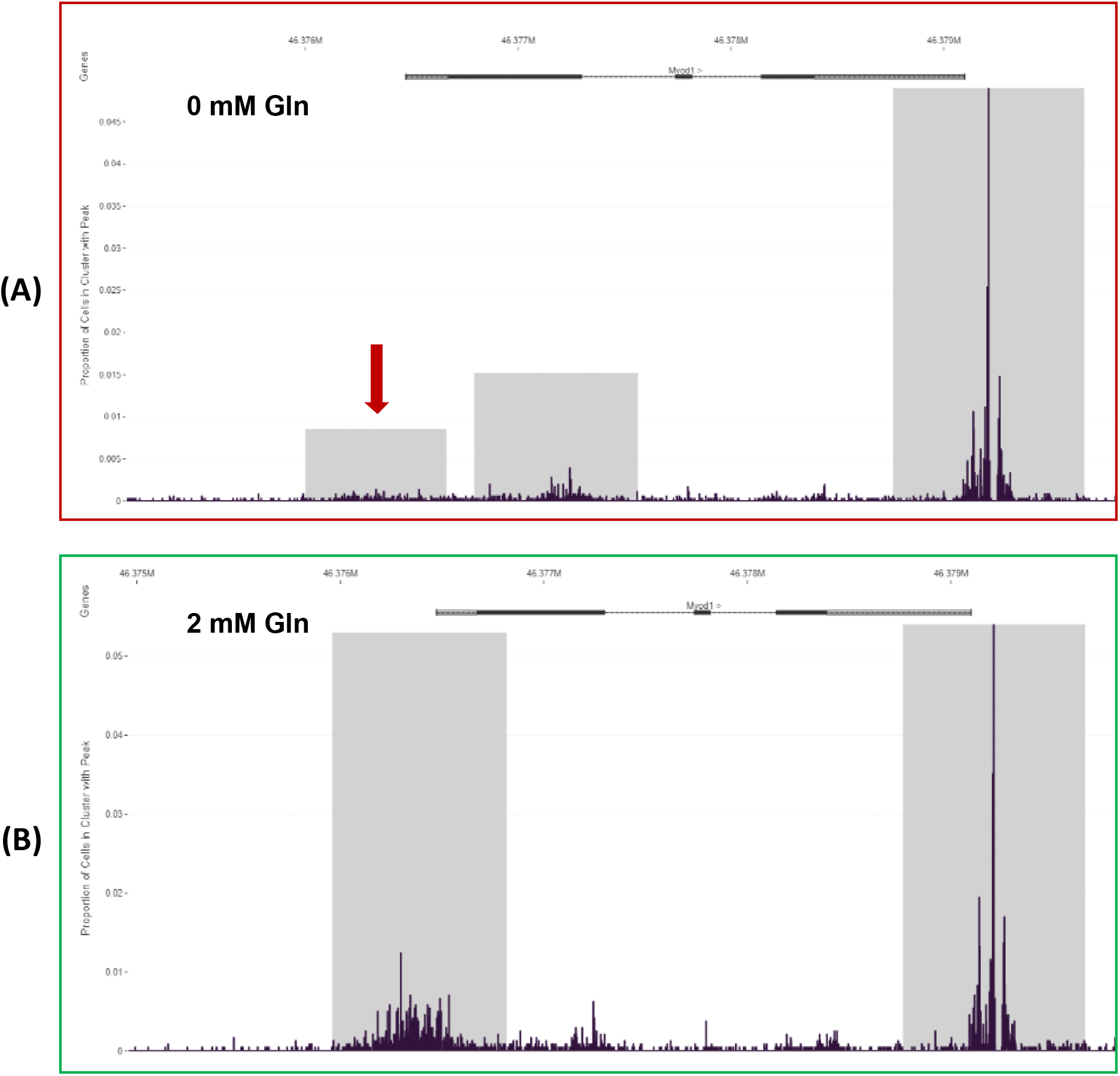
Differential chromatin accessibility of the myogenic regulatory gene MyoD1 under varying glutamine conditions. **(A)** Chromatin accessibility of the MyoD1 gene in cells treated with 0 mM Gln, visualized using the Loupe Browser. The MyoD1 locus shows reduced accessibility under Gln-depleted conditions. **(B)** Chromatin accessibility of the MyoD1 gene in cells treated with 2 mM Gln, visualized using the Loupe Browser. The MyoD1 locus shows increased accessibility under Gln-sufficient conditions.

## DISCUSSION

Our findings establish that Gln availability regulates MPC proliferation by linking extracellular nutrient status to intracellular metabolic rewiring and epigenetic remodeling (**Figure 7**). We observed that 2 mM Gln supports optimal MPC growth, whereas Gln deprivation restricts proliferation and induces G0/G1 cell cycle arrest. Importantly, this effect is reversible, indicating that Gln availability functions as a dynamic and modifiable signal for myogenic progression. Previous studies highlighted the role of Gln in protein synthesis and anabolic signaling^11,12^, our findings emphasize the necessity of intracellular Gln catabolism, vs protein synthesis *per se*, for cell growth^11^; inhibiting GLS, recapitulated the effects of Gln deprivation, confirming the necessity of mitochondrial Gln metabolism for MPC proliferation.

**Figure 7.**
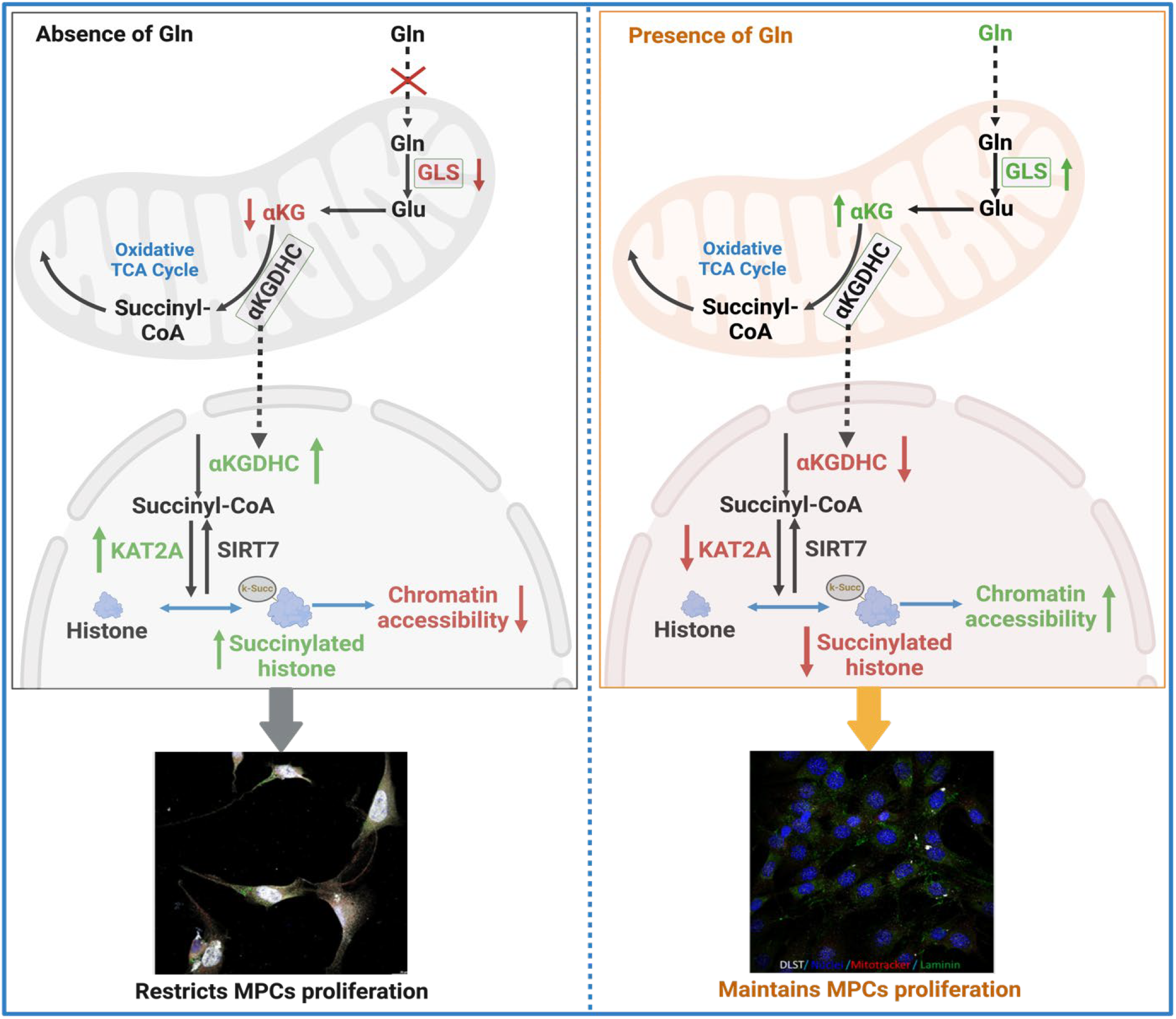
Schematic illustration of essentiality of extracellular Gln in promoting the proliferation of MPCs by enhancing chromatin accessibility of myogenic regulatory factor regions.

A central and novel finding of this study is that Gln deprivation induces nuclear translocation of mitochondrial α-KGDHC components-DLST and OGDH. This spatial redistribution of core mitochondrial enzymes suggests a reorganization of metabolic machinery in response to nutrient stress. This finding is consistent with emerging evidence in non-muscle tissues^15–17,29,32^, such translocation supports the idea that mitochondrial enzymes can adopt nuclear functions under stress conditions, including nutrient exposures. In the nucleus, α-KGDHC enzymes are known to interact with chromatin modifiers^29^. We observed a concurrent increase in the nuclear KAT2A, a succinyltransferase that utilizes succinyl-, from succinyl-CoA, to modify lysine residues, including those on histones. This nuclear accumulation of DLST, OGDH, and KAT2A, with no change in nuclear desuccinylase, SIRT7, suggests a shift toward nuclear (hyper)succinylation in the absence of extracellular Gln.

Importantly, this nuclear succinylation is not simply a secondary consequence of metabolic alterations but plays an active role in regulating chromatin accessibility. Single-nucleus ATAC-seq revealed decreased chromatin accessibility at key myogenic loci, including MyoD1, under Gln-deprived conditions. These findings support a model in which nuclear succinyl-CoA, generated by translocated α-KGDHC, acts as a glutamine-sensitive epigenetic signal that modifies the chromatin landscape. The increased succinylation of histones, such as H2B1B, H2AJ, and H3, likely perturbs nucleosome dynamics and restricts transcription factor access, leading to transcriptional repression of genes critical for MPC proliferation and lineage commitment. This aligns with previous studies demonstrating that histone succinylation is responsive to metabolic status and capable of modulating gene expression^29–31,33^.

Interestingly, while histone and structural protein succinylation increased under Gln deprivation, succinylation of TCA cycle enzymes decreased. This suggests a selective and compartmentalized redistribution of succinyl-CoA pools, reflecting a functional repurposing of metabolic intermediates toward nuclear processes. Similar patterns of succinyl-CoA utilization have been reported in hepatocytes and cancer cells under metabolic stress, including hypoxia, fasting, or oncogenic transformation^34–36^. Our findings extend this paradigm to muscle progenitor cell biology, suggesting a conserved response in which cells adaptively reallocate metabolic resources for epigenetic control. These results contribute to a growing understanding of how intermediary metabolism governs the epigenetic landscape. Succinyl-CoA, like other TCA-derived metabolites (e.g., acetyl-CoA), serves not only as an intermediate in energy production but also as a cofactor for chromatin-modifying enzymes^37,38^. Our study demonstrates that Gln availability orchestrates a mitochondrial–nuclear signaling axis that regulates histone succinylation and transcriptional accessibility at myogenic loci.

In summary, Gln functions as a metabolic nexus in MPCs, linking metabolism to nuclear epigenetic signaling. Extracellular Gln deprivation triggers the nuclear localization of mitochondrial enzymes-α-KGDHC, alters succinyl-CoA compartmentalization, and promotes widespread histone succinylation. These changes culminate in the repression of genes essential for MPC proliferation. This integrated mitochondrial–nuclear response provides a mechanistic framework for how extracellular nutrient availability is transduced into epigenetic and transcriptional outcomes, with important implications for skeletal muscle regeneration and perhaps metabolic disease.

## CONCLUSION

In conclusion, our study identified a novel role of extracellular Gln availability that alters chromatin accessibility and myogenic gene expression through metabolic and epigenetic coupling. We demonstrate that Gln availability regulates not only mitochondrial metabolism but also epigenetic remodeling via nuclear succinyl-CoA–mediated histone succinylation.

Upon extracellular Gln deprivation, mitochondrial enzymes such as α-KGDHC localize in the nucleus, where they drive succinylation of histones and restrict chromatin accessibility at key myogenic loci. This metabolic-epigenetic coupling results in transcriptional repression of genes essential for myogenic progression. These results highlight how metabolic flexibility and substrate availability determine the fate of muscle progenitors. Targeting this metabolic– epigenetic interface may offer new strategies to enhance muscle regeneration and mitigate muscle-wasting conditions associated with nutrient stress.

## DATA AVAILABILITY

All data generated related to the findings described in this paper are available upon request from the corresponding author.

