## Supplementary file for "Glutamine deficiency enhances nuclear localization of TCA cycle enzymes and epigenetic modifications, impairing myogenesis"

**Title: Glutamine deficiency enhances nuclear localization of mitochondrial-TCA cycle enzymes and impairs myogenesis**

**Supplementary files:** Three supplementary figures (S1-3) and three tables (Table S1-3).

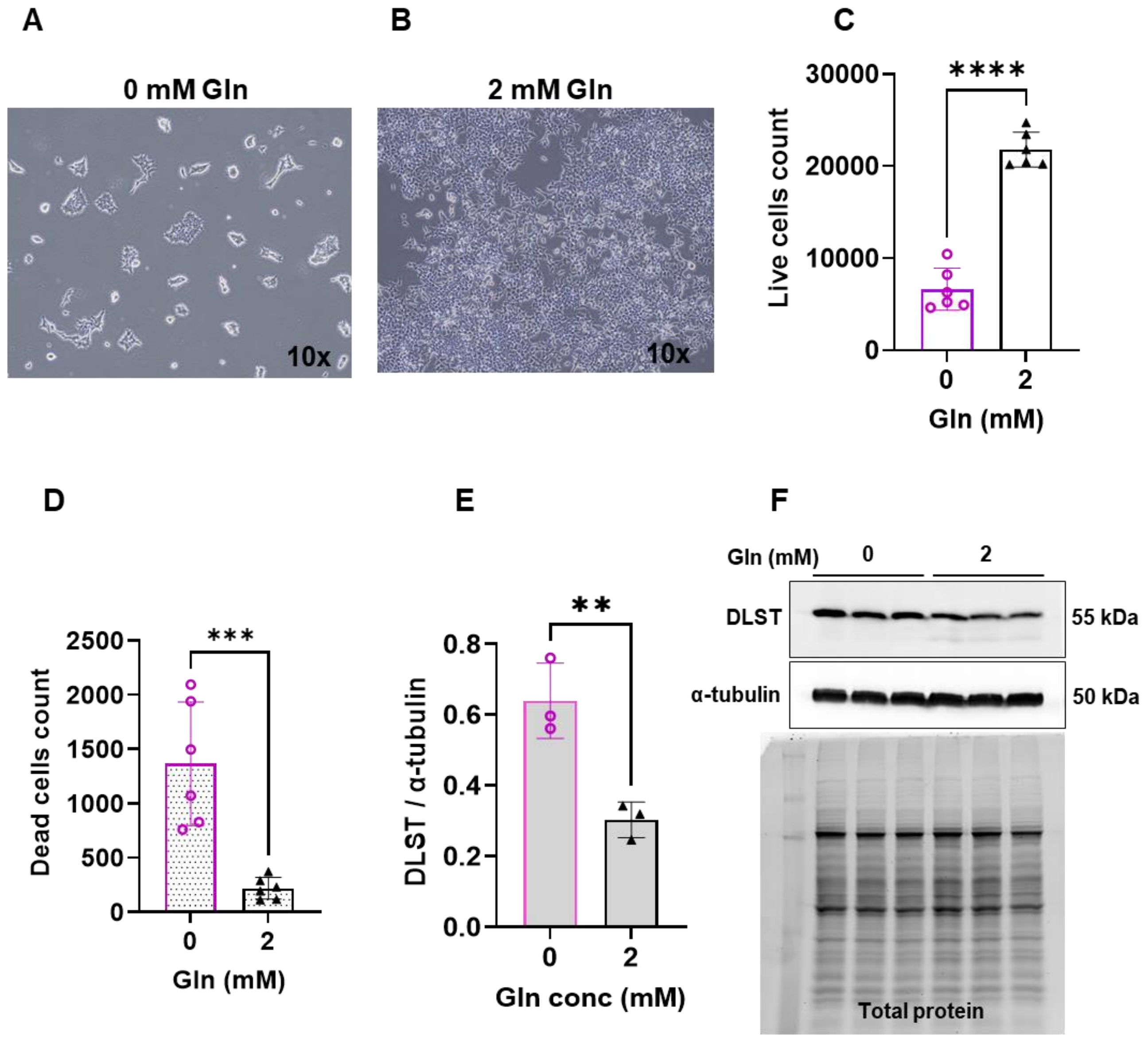


**Supplementary Figure 1. Extracellular Gln availability impacts HEK293 cell growth and DLST protein levels.**

**(A, B)** Representative brightfield images of HEK293 cells cultured for 96 hours in growth media without (0 mM) **(A)** and with (2 mM) Gln **(B)**. Images were acquired at 10× magnification.

**(C)** Quantification of live cell numbers shows a Gln-dependent increase in proliferation (n = 6).

**(D)** Cell death decreases with increasing extracellular Gln (n = 6).

**(E)** DLST protein levels are elevated under Gln-depleted conditions (n = 3).

**(F)** Representative immunoblot showing DLST, α-tubulin, and total protein levels.

All data were presented as mean ± SD. Level of significance p≤0.05; * **p≤0.01; ****p≤0.0001.

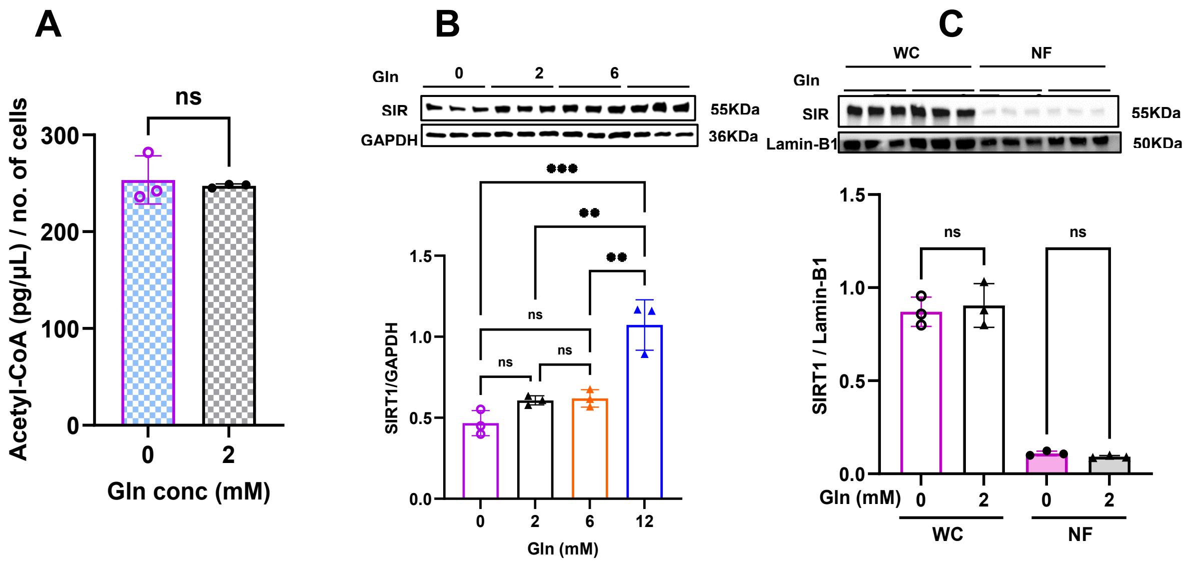


**Supplementary Figure 2. Effect of glutamine availability on Acetyl-CoA and SIRT1 level**.
**(A)** Acetyl-CoA unchanged with varying Gln availability, n=3.
**(B)** SIRT1 protein levels in whole-cell lysates with varying Gln availability, n=3.
**(C)** SIRT1 protein levels in the nuclear fraction unchanged with varying Gln levels, n=3.
All data were presented as mean ± SD. Level of significance p≤0.05; *p≤0.05; **p≤0.01; ****p≤0.0001. ns - non significant, WC - whole cell, NF - nuclear fraction.

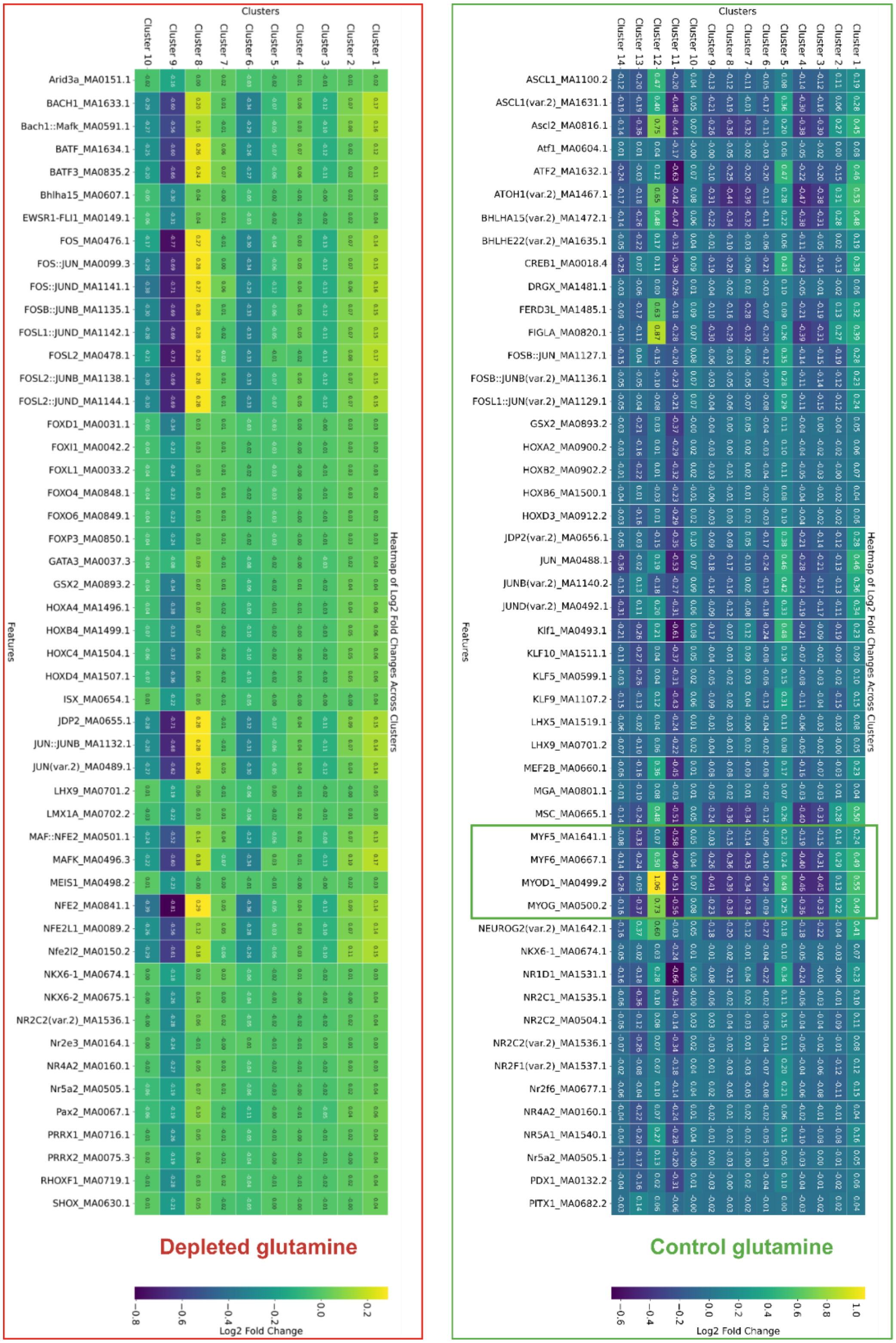


**Supplementary Figure 3. Significantly enriched motifs identified in glutamine depleted and control treated cells.** Motif enrichment analysis showed that motifs regions of myogenic regulatory factors were available for binding for transcription factors in case of control Gln condition (supplemented with 2 mM Gln).

**Table S1. Primers used in RT-qPCR experiments**

| Name | Forward primer | Reverse primer |
|---|---|---|
| MyoD (mus) | 5'-ACTTTCTGGAGCCCTCCTGGC-3' | 5'-TTTGTTGCACTACACAGCATG-3' |
| PAX7 (mus) | 5'-GACGACGAGGAAGGAGACAA-3 | 5'-ACATCTGAGCCCTCATCCAG-3' |
| β-actin (mus) | 5'-AAATCGTGCGTGACATCAA-3' | 5'-AAGGAAGGCTGGAAAAGAGC-3' |

**Table S2. Antibodies and their application**

| **Antibody name** | **Catalogue Number** | **Source** | **Dilution ratio** | | | |
|---|---|---|---|---|---|---|
| | | | *WB | **ICC | ***ICW | ****IP |
| Anti-MyoD antibody | sc-37740 | Santa Cruz | 1:1000 | | 1:100 | |
| Anti-MyoG antibody | sc-52903 | Santa Cruz | 1:500 | | | |
| Anti-PAX7 antibody | PA1-117 | Invitrogen | 1:500 | | 1:100 | |
| Anti-SLC1A5 antibody | PA5-50527 | Invitrogen | 1:1000 | | | |
| Anti-DLST antibody | PA5-52843 | Invitrogen | 1:1000 | 1:100 | | |
| Anti-OGDH antibody | ab307369 | Abcam | 1:1000 | | | |
| Anti-SIRT1 antibody | ab110304 | Abcam | 1:1000 | | | |
| Anti-SIRT7 antibody | ab259968 | Abcam | 1:1000 | | | |
| Anti-pan k-succinylation | PTM-419 | PTM BIO | 1:500 | | | 1:100 |
| Anti- KAT2A antibody | ab217876 | Abcam | 1:1000 | | | |
| Anti- α-tubulin antibody | 9099 | Cell Signaling | 1:1000 | | | |
| Anti-GAPDH antibody | 60004-1-1 | Protein Tech | 1:1000 | | | |
| Anti-Laminin antibody | L9393 | Sigma | | 1:500 | | |
| Anti-mouse IgG (tagged) | 7076s | Cell Signaling | 1:5000 | | | |
| Anti-rabbit IgG (tagged) | 7074s | Cell Signaling | 1:5000 | | | |
| Donkey Anti-Rabbit (Alexa Flour-647) | A31573 | Invitrogen | | 1:250 | | |
| Goat Anti-Mouse (Alexa Flour-488) | A32723TR | Invitrogen | | 1:250 | | |
| Donkey anti-mouse 647 | IRDye® 680RD | LI-COR | | | 1:800 | |
| Goat anti-rabbit | IRDye® 800CW | LI-COR | | | 1:800 | |
| Anti-mouse IgG (Untagged) | sc-2025 | Santa Cruz | | | | 1:100 |
| *WB-Western Blot; **ICC-Immunocytochemistry; ***ICW- In-Cell Western; ****IP-Immuno-precipitation; | | | | | | |

**Table S3. Total spectral counts of differential succinylated proteins**

| S.No. | RepID | 0mM Gln | 12mM Gln |
|---|---|---|---|
| 1 | MYH9 | 1162 | 555 |
| 2 | Q61276 | 157 | 58 |
| 3 | MYH10 | 156 | 67 |
| 4 | E9Q1F2 | 140 | 53 |
| 5 | Q61852 | 87 | 31 |
| 6 | ODO2 **(DLST)** | 58 | 148 |
| 7 | VIME | 43 | 13 |
| 8 | FINC | 40 | 3 |

| | | | |
|---|---|---|---|
| 9 | K2C5 | 38 | 24 |
| 10 | K1C10 | 34 | 12 |
| 11 | B2RTP7 | 32 | 15 |
| 12 | K2C1 | 28 | 19 |
| 13 | H2B1B | 26 | 0 |
| 14 | Q0VDR7 | 25 | 19 |
| 15 | H2AJ | 20 | 0 |
| 16 | K1C17 | 17 | 9 |
| 17 | E9Q0F0 | 15 | 13 |
| 18 | K1C42 | 15 | 0 |
| 19 | LMNA | 15 | 0 |
| 20 | IGHG1 | 14 | 13 |
| 21 | F8WI35 **(H3)** | 12 | 0 |
| 22 | IGHG3 | 12 | 10 |
| 23 | ALBU | 10 | 11 |
| 24 | HVM58 | 10 | 16 |
| 25 | MYL3 | 10 | 7 |
| 26 | Q8BP43 | 10 | 0 |
| 27 | B1AXB9 | 9 | 10 |
| 28 | KIF4 | 9 | 4 |
| 29 | RS27A | 9 | 0 |
| 30 | ODO1 **(OGDH)** | 8 | 22 |
| 31 | NPT2B | 7 | 21 |
| 32 | A2A4U6 | 5 | 2 |
| 33 | RL7 | 5 | 1 |
| 34 | F6SVV1 | 4 | 0 |
| 35 | PLEC | 4 | 1 |
| 36 | Q7M754 | 4 | 3 |
| 37 | TRI56 | 4 | 32 |
| 38 | XRCC6 | 4 | 1 |
| 39 | CAPZB | 3 | 0 |
| 40 | CDCA2 | 3 | 0 |
| 41 | DESM | 3 | 0 |
| 42 | H11 | 3 | 1 |
| 43 | LLPH | 3 | 0 |
| 44 | LRP2 | 3 | 1 |
| 45 | Q059P4 | 3 | 0 |
| 46 | RS17 | 3 | 0 |
| 47 | SCN9A | 3 | 0 |
| 48 | SYCP2 | 3 | 7 |
| 49 | TITIN | 3 | 8 |

| | | | |
|---|---|---|---|
| 50 | TMOD3 | 3 | 0 |
| 51 | B2RUK7 | 2 | 0 |
| 52 | B9EHN9 | 2 | 2 |
| 53 | B9EKS2 | 2 | 0 |
| 54 | GNL3 | 2 | 0 |
| 55 | HERC2 | 2 | 0 |
| 56 | IGF1R | 2 | 0 |
| 57 | J3QM72 | 2 | 0 |
| 58 | NEK10 | 2 | 0 |
| 59 | ODP2 | 2 | 2 |
| 60 | PCKGM | 2 | 5 |
| 61 | PGBM | 2 | 0 |
| 62 | Q8BIV1 | 2 | 0 |
| 63 | Q99L75 | 2 | 0 |
| 64 | SSUH2 | 2 | 1 |
| 65 | TDRD1 | 2 | 1 |
| 66 | V9GWY0 | 2 | 0 |
| 67 | VP26A | 2 | 0 |
| 68 | ZFY2 | 2 | 1 |
| 69 | ZMYM4 | 2 | 0 |
| 70 | MT2 | 1 | 2 |
| 71 | NTKL | 1 | 2 |
| 72 | TNR6A | 1 | 3 |
| 73 | ABCA4 | 0 | 2 |
| 74 | CHRD | 0 | 3 |
| 75 | E9Q7N9 | 0 | 2 |
| 76 | FBN2 | 0 | 2 |
| 77 | FIL1L | 0 | 2 |
| 78 | HS90B | 0 | 2 |
| 79 | LNEBL | 0 | 2 |
| 80 | O35452 | 0 | 2 |
| 81 | O88840 | 0 | 2 |
| 82 | OTOGL | 0 | 2 |
| 83 | SV421 | 0 | 2 |
| 84 | TES | 0 | 2 |
| 85 | UBR1 | 0 | 2 |
| 86 | WDR3 | 0 | 2 |
| 87 | WFC6A | 0 | 3 |
